# Continuous thermal sensitivity of gene expression following acclimation in *Drosophila subobscura*

**DOI:** 10.64898/2026.08.05.743044

**Authors:** Ellie S. Tushar, Mara Heilig, Adam Haddad, James A. deMayo, Gregory J. Ragland

## Abstract

The physiology of ectotherms can change substantially during acclimation to changing environmental temperature. The role of transcription in acclimation responses has been well-established, but it remains unclear whether transcriptional regulation generally reflects abrupt changes after surpassing temperature thresholds, or whether transcript abundance is a relatively monotonic, continuous function of acclimation temperature. In this study we exposed adult male *Drosophila subobscura* flies to four different 96-hour acclimation treatments at temperatures that were not acutely stressful but ranged from relatively cold (10°C) to relatively warm (27°C) with respect to standard rearing conditions. Transcriptome sequencing of whole-body homogenates (mRNAseq) revealed a massive, transcriptome-wide response across acclimation temperatures, with a marked overrepresentation of genes that were continuously and monotonically up- and down-regulated in response to increasing acclimation temperature. Though some genes showed more complex relationships consistent with putative threshold responses, a high percentage of the differentially expressed transcriptome (42%) showed continuous and strictly monotonic relationships. Functional enrichment suggested continuous up-regulation of spermatogenesis-related transcripts with increasing temperature and continuous up-regulation of oxidative phosphorylation-related transcripts with decreasing temperature, illustrating contrasting patterns consistent with previous studies of thermal sensitivity of male reproduction and metabolic compensation in the cold. Thus, continuous thermal sensitivity of transcription is a hallmark of acclimation in *D. subobscura* that likely underlies the continuous thermal sensitivity of downstream physiological processes. We also provide evidence for shared transcriptomic responses across short-term acclimation (this study) vs. published results for long-term, developmental acclimation.

## 1. Introduction

Physiological adjustments during thermal acclimation play an important role in determining survival (Levins, 1969; Narum et al., 2013), reproduction (Porcelli et al., 2017; Wilson et al., 2007), and ultimately fitness and persistence in variable environments (Leroi et al., 1994) (Morley et al., 2019). Physiological acclimation is particularly well described for ectothermic animals that have long served as models for comparative physiology of thermal performance. By the 1960’s comparative physiologists had identified a key role of alternate protein isoforms of metabolic enzymes in the acclimation process (Somero, 2004). These results in turn implied a potential role of transcriptional regulation that feeds forward to protein/isoform abundances. The advent of cDNA microarrays and later high throughput sequencing corroborated this hypothesis, suggesting that acclimation at different temperatures affects many physiological processes through widespread changes in gene expression (Franke et al., 2019; MacMillan et al., 2016; Podrabsky and Somero, 2004; Stillman and Tagmount, 2009).

Though transcription clearly plays a role in acclimation, quantitative relationships between transcription and acclimation temperature (reaction norms) have rarely been investigated. Often, transcriptomic studies test for acclimation to a relatively cold or relatively warm experimental temperature compared to an organism’s presumed ‘normal’, if not optimal range of temperatures (Bay and Palumbi, 2015; Drown et al., 2022; Liu et al., 2017; Oomen and Hutchings, 2017; Windisch et al., 2014). Acclimation responses observed in these studies could involve transcriptional changes that only occur after surpassing threshold warm or cold temperatures (e.g., the classic heat shock protein response; Feder and Hofmann, 1999). Alternatively, transcriptional responses to acclimation may reflect relatively continuous thermal sensitivity of gene expression yielding linear or curvilinear relationships with temperature, as has been demonstrated in long-term, egg-to-adult acclimation in the *Drosophila melanogaster* flies (Chen et al., 2015). These hypotheses are not mutually exclusive in the sense that different genes may exhibit different expression patterns. Thus, the most relevant question is whether certain patterns (i.e., reaction norm shapes) play a dominant role in the transcriptome-wide response to acclimation temperature.

Here, we test for overrepresentation of certain reaction norm shapes and evaluate the relative transcriptomic contribution of continuous, monotonic reaction norms to short-term, adult acclimation in *Drosophila subobscura*. We also briefly explore likely physiological processes regulated by transcriptional acclimation and similarities between short-term acclimation in adult flies (this study) and egg-to-adult, long-term acclimation (Chen et al., 2015).

## 2. Methods

### 2.1 Experimental Design

We performed experiments on an isofemale line (US National Drosophila Species Stock Center at Cornell, sku 14011-0131.16, collected from East Sussex, UK, ∼ 51°N, 0.3°E) of *Drosophila subobscura*, a Palearctic *Drosophila* fly that has recently colonized North and South America (Prevosti et al., 1988). We designed our experiment to capture gene expression following adult acclimation at one high and two low temperatures relative to a temperature near the fitness optimum (∼22° C; MacLean et al., 2019), yielding four acclimation temperatures: 10, 15, 21, and 27° C. We chose these temperatures based on prior thermal sensitivity studies showing that Chilean lines (∼ 41° S) successfully reproduce at constant temperatures from 13 - 22° C (Laayouni et al., 2007), and Danish lines (56.4°N) show a fitness optimum near 22°C (MacLean et al., 2019). We allowed experimental flies to acclimate for four days, similar to acclimation durations shown to elicit a robust transcriptional response in *D. melanogaster* (MacMillan et al., 2016).

### 2.2 Fly Rearing and Experimental Protocol

All fly stocks used to produce experimental flies were reared at 21°C with 50% relative humidity (RH) and a 14:10 photoperiod following (Vidrio et al., 2026), with full details of rearing and experimental handling in Supplemental Methods. Newly eclosed males were sorted into fresh vials and placed in incubators set to either 10, 15, 21, or 27° C with the same RH and photoperiod as above. Following a four-day acclimation treatment, flies were aspirated into empty vials, flash frozen, and stored at -80°C. Frozen flies were haphazardly sorted into treatment pools of 5 flies, with each pool representing a roughly random sample across eclosion times.

### 2.3 RNA Preparation, Sequencing, and Informatics

Total RNA was extracted from four pools of 5 male flies (4 biological replicates) per acclimation treatment and sequenced to a target depth of 100 million 150bp paired-end reads per sample. Initial quality was assessed with FastQC (Andrews, n.d.), then reads were aligned to the NCBI *D. subobscura* reference genome (GCF_008121235.1) using STAR (Dobin et al., 2013), retaining uniquely mapping reads to generate counts per gene. For full details see supplemental methods and Zenodo archive.

### 2.4 Data Analysis

All analyses were performed using standalone software or packages in R version 4.5.2 (R Core Team, 2025), and detailed explanations of all steps are available in supplemental methods. After initial inspection of multidimensional scaling (MDS) clustering (Fig. S1) and further inspection of female-specific gene expression Table S1) we removed one female-contaminated sample, leaving 3 replicates in the 15° C treatment. Subsequent MDS analysis revealed non-overlapping clusters for each temperature treatment (Fig. S2).

#### 2.4.1 Fitting a discrete model: temperature as a discrete predictor

We fit a discrete generalized linear model (glm) treating temperature as a factor, estimating log_2_ fold changes from contrasts of all temperatures back to 10°C, and calculating Benjamini Hochberg-corrected p-values (FDR). We designated a subset of ‘discrete plastic’ genes as the subset with at least one contrast yielding an FDR value < 0.05, evidence of differential expression.

#### 2.4.2 Fitting a continuous model: temperature as a continuous predictor

We also fit a continuous glm treating temperature as a continuous predictor and including an intercept, a linear term, and a quadratic term. The resulting linear and quadratic coefficients represent the estimated linear and quadratic change expression across temperatures. We then designated a subset of ‘continuous plastic’ genes as those with FDR < 0.05 associated with the linear or the quadratic term.

#### 2.4.3 Clustering of ‘discrete plastic’ genes and functional enrichment of discrete clusters

We used the ‘discrete plastic’ set of genes as input for the short time series expression miner (STEM) algorithm (Ernst et al., 2005; Ernst and Bar-Joseph, 2006) to identify ‘template’ profiles of thermal reaction norms that were statistically overrepresented in the data (temperature serves as the ‘time’ variable). This approach clusters genes into template shapes, or profiles (here, reaction norms) using Pearson correlation (*r*) and uses permutation and FDR correction to assign a corrected p-value for each cluster (Table S2) We combined similar profiles (*r* > 0.7) into clusters, and we report trajectories (in log_2_ fold change vs. 10°C) for each gene in each profile in Table S3. We then conducted gene enrichment analysis on each cluster against the Gene Ontology biological processes (GO:BP) and Kyoto Encyclopedia of Genes and Genomes pathways (KEGG) databases using *gprofiler2* with default settings against *D. melanogaster* annotations (Flybase release 6.63) based on Blastp best hits (Table S4-S22).

After initial inspection suggested an excess of monotonic relationships with temperature, we also tested whether strictly increasing or decreasing relationships (all estimated log fold changes relative to 10°C > 0 or < 0) were overrepresented in the full set of gene-wise expression trajectories. We calculated the percent of discrete plastic genes either strictly increasing or decreasing and compared against estimates from 1000 random permutations of the temperature order to calculate a p-value from an empirical null distribution.

#### 2.4.4 Clustering of ‘continuous plastic’ genes and comparison with developmental acclimation in D. melanogaster

In a second clustering approach, we used an analogous strategy to Chen et al. (2015) to cluster continuous plastic genes based on evidence for linear and quadratic polynomial regression terms. We clustered genes into the following classes: 1) genes with only a significant linear term (FDR < 0.05) as increasing or decreasing according to the sign of the linear coefficient; 2) genes with a significant quadratic term (FDR<0.05) as “concave-up” or “concave-down” according to the sign of the quadratic coefficient and increasing or decreasing according to the sign of the linear coefficient; 3) a subset of genes with a significant quadratic term as U-shaped or bell-shaped if the diference in mean expression between the temperature extremes (10°C and 27°C) was less than 80% of the total range in mean expression across all four temperatures (following Chen et al, 2015). This complimentary approach produces a simpler set of clusters (compared to STEM) that identify a smaller subset of straight line and curvilinear relationships with temperature.

Using this approach also allowed us to directly compare our results to those of Chen et al. (2015). Because Chen et al. (2015) do not report (unfitted) individual expression trajectories for each gene, we instead asked whether lists of genes assigned to a particular cluster (e.g., linear, increasing) in our study were enriched (fisher exact test; odds ratio > 1; p < 0.05) for the list of genes from *D. melanogaster* grouped in the comparable cluster in Chen et al. (2015).

## 3. Results

### 3.1 Most of the transcriptome responds to acclimation at one or more tested temperatures

Discrete linear models identified a massive transcriptional response to acclimation temperature, with 9,272 genes (discrete plastic genes) differentially expressed in at least one contrast back to 10°C, representing 76% of genes expressed at levels exceeding our filtering threshold (see methods).

### 3.2 A high proportion of plastically expressed genes are monotonically related to temperature

Clustering of expression acclimation patterns with STEM revealed marked enrichment for genes that monotonically increase or decrease across acclimation temperatures (Fig. 1). The four statistically overrepresented clusters (shapes) with the most genes (Clusters I. – IV., 36%, or 3,315 out of 9,208 discrete plastic genes) represented exclusively weakly or strictly increasing/decreasing profiles. Further, 64 % of genes in the 5 clusters containing the highest numbers of genes (Clusters I. – V.) were strictly monotonic (Table S2). There were non-monotonic, overrepresented reaction norm shapes, but all were relatively simple relationships with a single peak in expression (e.g., Clusters VII., IX.).

**Figure 1.**
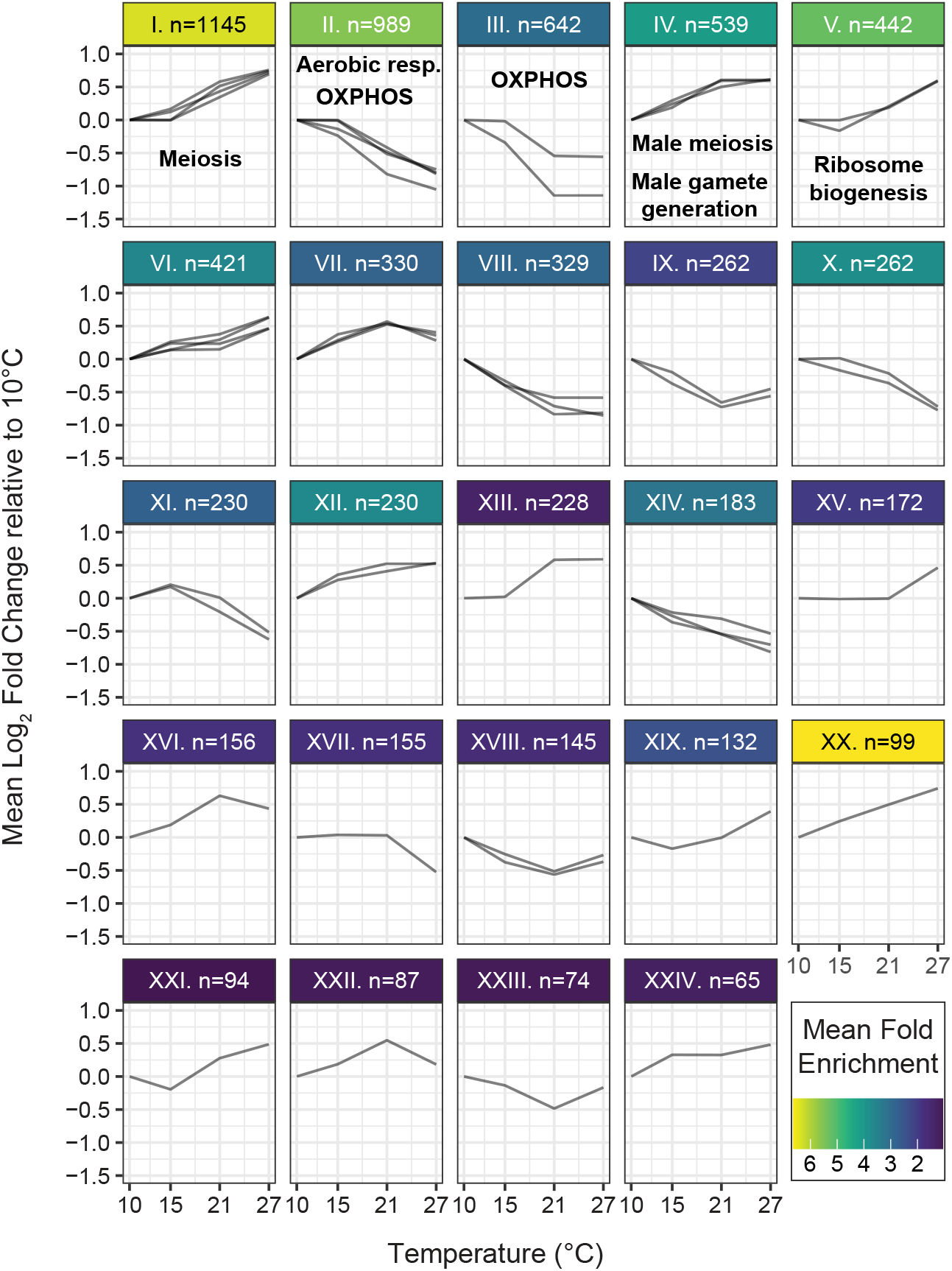
Short time series expression profiles. Each line depicts the mean expression trajectory (differential expression relative to expression at 10°C) for genes mapped into a single profile. Each panel represents a cluster overrepresented in the data (FDR < 0.05), and clusters with multiple lines include profiles with pairwise Pearson correlations > 0.7. Panels include the total number of genes per cluster (n) and color indicating the magnitude of the mean fold enrichment of the member profiles (number of genes in a profile / number expected from randomly permuted data). Select, enriched functional categories are also indicated for the five largest clusters, see Results section for explanation. Category names abbreviated for presentation, full names and database origins are as follows: meiotic cell cycle (GO:BP, cluster I.), aerobic respiration and oxidative phosphorylation (GO:BP, cluster II.), oxidative phosphorylation (KEGG, cluster III.), male meiosis I and male gamete generation (GO:BP, cluster IV.), and ribosome biogenesis (GO:BP, cluster V.).

Recognizing that not all member genes in a cluster exactly follow the template shape, we also conducted gene-wise calculations and a permutation test showing that 42% of discrete plastic genes had strictly increasing/decreasing trajectories (log fold changes relative to 10°C), 3.4 fold greater than expected by chance (p < 0.001).

Applying the continuous linear modeling approach also revealed clusters consistent with monotonic relationships. Indeed, the largest clusters of continuous plastic genes followed straight-line relationships with temperature (linear term with FDR < 0.05, quadratic term with FDR > 0.05; clusters a and b, Fig. 2).

**Figure 2.**
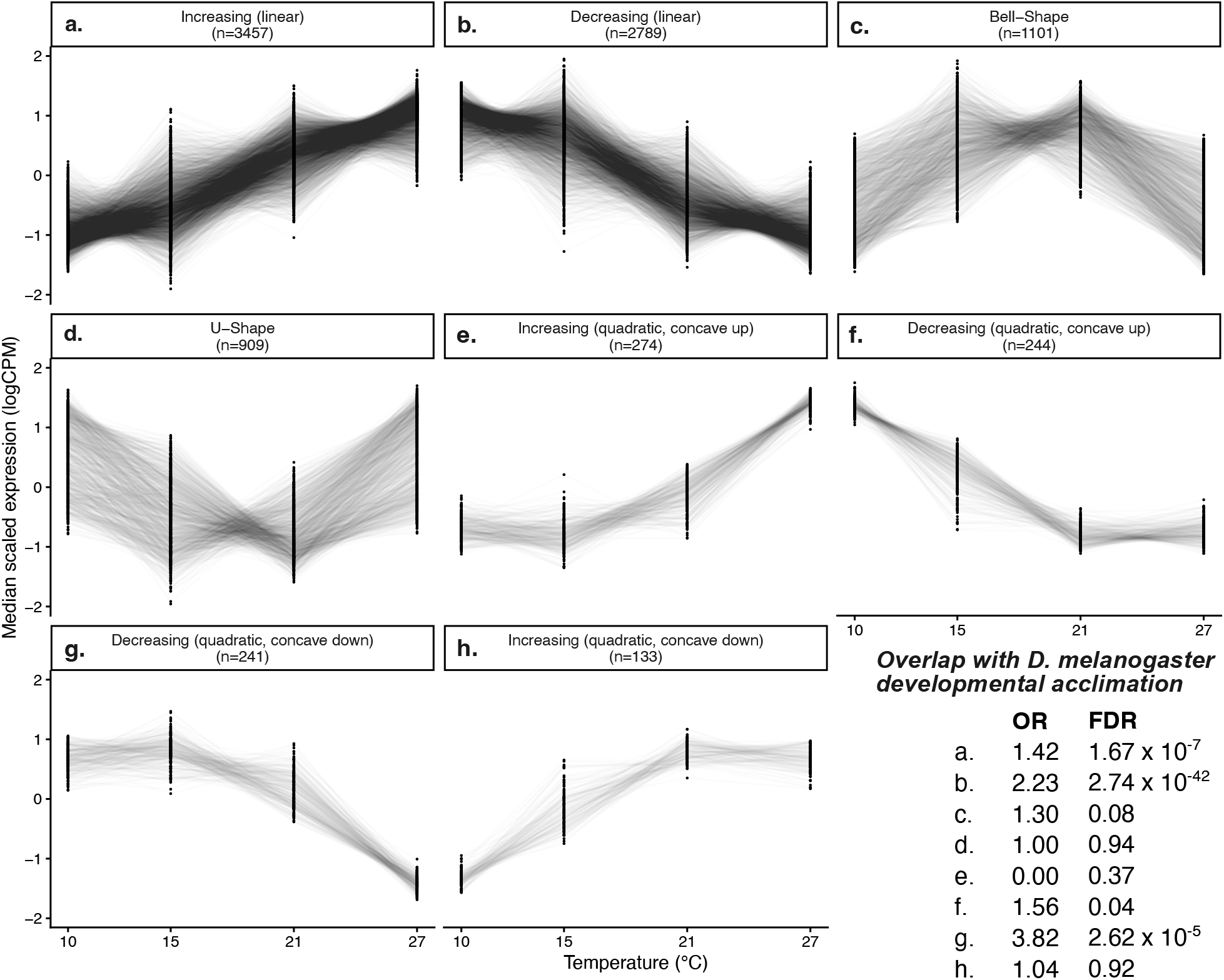
Clustering of continuously plastic genes and comparison with developmental acclimation in *Drosophila melanogaster* (Chen et al. 2015). Normalized read counts were standardized to a mean value of 1 for illustration purposes; median values for each replicate at each temperature are presented as black dots. Genes with significant linear (a,b) or quadratic (c-h) relationships with acclimation temperature were assigned to clusters based on the sign and significance of their regression coefficients. Panel headers indicate the cluster class as described in the methods and the number of member genes (n). Overlap between clusters from this study and similar clusters reported for developmental acclimation in *D. melanogaster* by Chen et al. (2015) was evaluated for enrichment using Fisher’s exact tests. The resulting odds ratios (ORs) and FDR corrected *p*-values are presented in the lower right panel.

Altogether, these results suggest that expression of a large complement (though not the majority) of the plastic transcriptome is consistently and monotonically related to temperature over the measured range. Though there were certainly additional, more complex patterns of expression (e.g., non-monotonic profiles also overrepresented in the STEM clustering), relatively few genes were up- or down-regulated between only two measured acclimation temperatures (e.g., clusters XIII., XV., and XVII).

### 3.3 The plastic response reflects upregulation of male meiosis and gamete generation and downregulation of aerobic respiration with increasing temperature

An examination of enriched functional categories for the 5 most inclusive STEM clusters (I. – V.) reveals two clear functional trends. First, genes involved in meiosis (and in particular, male meiosis I), male gamete formation, and ribosome biogenesis were highly enriched in clusters associated with increased expression at higher temperatures (clusters I., IV., and V.). Second, genes involved in aerobic respiration and specifically oxidative phosphorylation were highly enriched in clusters associated with increased expression at low temperatures (clusters II. and III.). These patterns are consistent with relatively monotonic thermal sensitivity of transcription-level regulation of both processes, but in opposite directions. Here we highlight these particular functional categories 1) because of the strength of enrichment (all FDR < 0.03, most FDR < 1E-5) and/or the consistency in broad functional groupings across similar-shaped clusters, and 2) to highlight two highly plausible sets of processes as examples of continuous, largely monotonic thermal sensitivity (see discussion). For full enrichment results and statistics see Supplemental TableS4-S21.

### 3.4 Short term acclimation in D. subobscura adults induces similar responses compared to developmental acclimation D. melanogaster

We found robust evidence for similarities in expression patterns between our study and Chen et al. (2015) revealed by the results of the continuous clustering analysis (Fig. 2).

Specifically, all clusters of genes from our study with a net (and largely monotonic) decrease in expression with increasing temperature were enriched (FDR < 0.05) for genes that were assigned to the analogous cluster (e.g., linear, increasing in both studies) in Chen et al. (2015) (Fig. 2; clusters a, b, f, and g). We also found significant overlap with the Chen et al. (2015) study for the cluster of genes with a straight line, positive relationship with temperature (cluster a). As noted in the methods, these similarities were identified despite differences in species and sex between studies (male in our study, female in Chen et al., (2015)).

## 4. Discussion

### 4.1 Continuous thermal sensitivity of expression influences acclimation physiology

Acclimation induces a massive, transcriptome-wide, plastic response in *D. subobscura*, and our results suggest a preponderance of plastically-expressed genes that continuously, and often monotonically change expression across acclimation temperatures. In fact, we have likely underestimated the number of differentially expressed genes because whole-body homogenates surely mask tissue-specific thermal responses (e.g., Heilig et al., 2026). The transcriptome-wide scale of the acclimation response is consistent with similar results in other ectotherms (Franke et al., 2019; MacMillan et al., 2016; Podrabsky and Somero, 2004; Stillman and Tagmount, 2009), further supporting the prominent role of gene regulation during acclimation. Relatively continuous expression variation with acclimation temperature is also consistent with a previous study of egg-to-adult acclimation in *D. melanogaster* (Chen et al., 2015). These results would suggest that many physiological processes influenced by gene expression should also vary relatively continuously, and monotonically with acclimation temperature. Indeed, whole-organism traits such as the critical thermal maximum/minimum scale linearly with acclimation temperatures from 12 – 32°C in *D. melanogaster*, though metabolite profiles may exhibit curvilinear relationships (Schou et al., 2017). There are certainly examples of steep temperature thresholds for trait induction in ectotherms – heat shock protein (HSP) expression after surpassing a high temperature threshold is perhaps the most recognizable and best studied (Feder and Hofmann, 1999). However, these threshold responses may be induced by shorter exposure durations (e.g., threshold responses during rapid cold hardening over a few hours; Vidrio et al., 2026). Or, thresholds may manifest primarily when temperatures stray into stress-inducing extremes as is the case for HSP induction. In general, the effects of acclimation, and thus the shape of acclimation reaction norms, varies widely across traits, species, range of acclimation temperatures, and duration of the acclimation period (Colinet and Hoffmann, 2012; Rako and Hoffmann, 2006; Van Heerwaarden et al., 2024). Nevertheless, the available evidence suggests that continuous thermal sensitivity of gene expression governs acclimation across a relatively broad range of temperatures in *Drosophila* flies and possibly other ectotherms.

Functional enrichment suggests that these continuous expression profiles contribute to several processes that compensate for low metabolism at low temperatures and increase reproductive investment at higher temperatures. Genes associated with aerobic respiration were continuously upregulated with decreasing acclimation temperature in our experiments. These results are consistent with observations that *D. melanogaster* increases aerobic metabolism and ATP synthesis when acclimated at lower temperatures, hypothesized to reflect metabolic compensation to enhance cold tolerance and counteract Arrhenius effects that decrease metabolism in the cold (Berrigan, 1997; Colinet et al., 2017). In contrast, genes associated with spermatogenesis (male meiosis, male gamete generation) were continuously upregulated with increasing temperature. This trend is highly consistent with several studies showing that male fertility (Chakir et al., 2002; Colinet et al., 2025) and sperm production (Gandara and Drummond-Barbosa, 2023) increase with temperature in *D. melanogaster*. Moreover, increasing transcription of genes involved in ribosome biogenesis in our study is also consistent with predicted high translational activity in the post-meiotic stages of spermatogenesis (Fabian and Brill, 2012). Overall, these examples illustrate how continuous patterns of thermal sensitivity of transcription may underpin physiological processes with different and even contrasting thermal sensitivities.

### 4.2 Transcriptional similarities between short- and long-term acclimation

We identified similar gene clustering patterns across temperatures between our study (adult acclimation) and the study of egg to adult acclimation in *D. melanogaster* reported in Chen et al., (2015). This result implies conserved transcriptional acclimation across species (*D. subobscura* vs. *D. melanogaster*) and across sexes (male in this study, female in (Chen et al., 2015). Furthermore, enrichment of shared genes in four of these clusters strongly suggests functional similarities between short-term adult acclimation (this study) and long-term egg-to-adult acclimation (Chen et al., 2015). One possibility is that these similarities reflect a shared transcriptional response to acclimation regardless of life stage and duration of exposure. An alternative, and not mutually exclusive possibility is that adult acclimation dominates the egg-to-adult response such that developmental studies like Chen et al. (2015) largely capture recent, adult-specific plasticity rather than plastic expression due to the thermal experience of earlier developmental stages. Indeed, adult acclimation in *D. melanogaster* has been shown to reverse the effects of developmental acclimation at earlier stages (Angilletta et al., 2019). Moreover, Chen et al. (2015) sampled flies at 3-days post eclosion, and both *D. subobscura* (this study) and *D. melanogaster* (MacMillan et al., 2016) exhibit robust transcriptional responses following acclimation for four or six days, respectively. Thus, it seems likely that common responses to adult acclimation contribute to the observed overlap with results reported in Chen et al. (2015). Whether short-term and longer-term, developmental acclimation rely on shared or largely distinct molecular mechanisms remains an unresolved question, with existing evidence supporting both partial overlap (Metzger and Schulte 2018) and largely independent genetic architecture (Gerken et al., 2015). Further experiments that manipulate the duration and life stage of thermal acclimation could help to disentangle acclimation effects spanning different developmental time windows.

### 4.3 Conclusion

Continuous thermal sensitivity characterizes the expression pattern of thousands of genes during short-term (days), and possibly long-term (whole life cycle) acclimation in *Drosophila* flies. In turn, we predict that thermal sensitivity of expression drives the continuous thermal sensitivity of many higher-level processes and phenotypes altered during acclimation.

