## Supplemental Methods and Figures for "Continuous thermal sensitivity of gene expression following acclimation in *Drosophila subobscura*"

All code used to implement informatic and statistical procedures will available in a Zenodo archive upon acceptance of this manuscript.

2.2 Fly Rearing and Experimental Protocol

Progenitor experimental flies were initially reared in bottles at 21°C with 50% relative humidity and a 14:10 photoperiod. Vials containing 5 males and 5 females were then established from newly eclosed flies sampled from the bottles to standardize larval densities. Vials were cleared once larval activity was observed, and we monitored daily for new adult emergence once pupae started to form. On the day of adult emergence, males were sorted with light CO2 anesthesia and transferred into fresh vials at a maximum density of 10 flies/vial. Vials were then placed in one of four incubators (Percival DR-36VL incubator, Percival Scientific, Perry, IA) programmed with 14:10 photoperiod, 50% humidity, and either 10, 15, 21, and 27° C constant temperature. All rearing and experimental vials/bottles contained a cornmeal-agar-yeast diet described in (Vidrio et al., 2026). Following the four-day acclimation treatment, male flies were aspirated into empty vials to ensure no debris or dead flies were transferred into the sample. They then were tipped into a 15ml tube, flash frozen with liquid nitrogen, and immediately placed in a -80°C freezer to limit RNA degradation.

Once all male flies for an acclimation treatment were frozen, we haphazardly transferred individuals from different collection dates (variable due to variation in emergence timing) into microfuge tubes placed on dry ice until all tubes contained a total of 5 male flies. This ensured each biological replicate of 5 flies was randomized across emergence dates.

2.3 RNA Preparation, Sequencing, and Informatics

Total RNA was extracted from four-day old adult males using the Zymo Direct-zol RNA MiniPrep kit (Cat. R2050) following the manufacturer's protocol. Immediately following each batch of extractions, RNA concentration and purity were assessed using both a Qubit 4 fluorometer (Invitrogen; RNA Broad Range assay) and a Nanodrop One spectrophotometer (ThermoFisher Scientific). Eluted RNA was stored at –80°C until library preparation. All steps for library preparation and sequencing were performed by the University of Colorado Denver Anschutz Medical Campus Genomics Shared Resource facility. RNA purity and quantity were confirmed on a NanoDrop and integrity was determined with a TapeStation 4200 (Agilent) analysis prior to RNA-seq library preparation. The Universal Plus mRNA-Seq library preparation kit with NuQuant (Tecan) with an input of 200ng of total RNA was used to generate RNA-Seq libraries. Paired-end sequencing reads of 150bp were generated on a NovaSeq X Plus (Illumina) sequencer at a target depth of 100 million paired-end reads per sample. To achieve the target depth, each sample was multiplexed with other samples (not included in this study), and the larger pool of libraries was sequenced across four lanes. Raw sequencing reads were de-multiplexed using bcl2fastq and counts per gene were summed across all four lanes.

#### 2.4 Data Analysis

We initially assessed variation within and among samples using the plotMDS function (default settings including the top 500 most variable genes) from the limma package (Ritchie et al., 2015). We identified non-overlapping clusters for each temperature treatment, as well as one sample from the 15° C treatment that appeared as an outlier on the second dimension (Fig. S1). To investigate this outlier, we identified genes with a log fold-change >4 relative to other samples in the same treatment and used BLASTp to identify Drosophila melanogaster orthologs. These orthologs were analyzed for functional enrichment using DAVID, revealing clustered enrichment for processes related to female development and oogenesis. We examined normalized read counts for three genes (Table S1) with female-biased expression and found substantially elevated expression in the outlier sample relative to all other samples, regardless of treatment, consistent with contamination with one or more female flies. After removing these outliers, MDS plotting revealed non-overlapping, distinct cluster of samples for each temperature treatment. We used this reduced set of samples for all subsequent analyses.

We applied analyses that focus on identifying transcriptome-wide patterns rather than identifying individual candidate genes. Thus, we used relatively standard, but not overly conservative statistical criteria for identifying differentially expressed genes followed by clustering approaches that emphasize multi-gene patterns.

We used linear models with empirical Bayes moderation fitted with limma-trend (Ritchie et al., 2015) to test for differential expression across temperatures, filtering down to 12,267 genes with >= 10 mapped reads in > 95% of samples and using log(counts per million) as input. We fit two distinct models treating temperature as either a discrete or continuous predictor.

##### 2.4.1 *Fitting a discrete model: temperature as a discrete predictor* through 2.4.2 *Fitting a continuous model: temperature as a continuous predictor*

We fit the discrete glm with a four-parameter design matrix (intercept plus 3 temperature parameters) and then fit contrasts of each acclimation treatment back to the lowest temperature (10°C) to estimate $\log_2$ fold changes (relative to 10°C) and associated, Benjamini Hochberg-corrected p-values (FDR). We fit the linear glm with a three-parameter design matrix including an intercept, a linear term, and a quadratic term. The linear term was the acclimation temperature mean-centered across samples, and the quadratic term was the square of this centered linear term.

*2.5.3 Clustering of 'discrete plastic' genes and functional enrichment of discrete clusters*

Permutation test for monotonicity

Initial inspection of the STEM results suggested that a high proportion of genes clustered into profiles that were at least weakly monotonic (all profile changes relative to 10°C >= 0 or <= 0) over the entire range of acclimation temperatures, suggesting relatively consistent thermal sensitivity. We further explored this result by testing whether strictly increasing or decreasing relationships (all estimated log fold changes relative to 10°C > 0 or < 0) were overrepresented in the full set of gene-wise expression trajectories. We calculated the percent of discrete plastic genes either strictly increasing or decreasing and compared the estimated percentage against estimates from 1000 random permutations of the temperature order (independently permuting for each gene in each iteration) to calculate a p-value from an empirical null distribution.

2.4.4 *Clustering of 'continuous plastic' genes and comparison with developmental acclimation in D. melanogaster*

We used the best Blastp hits (*D. subobscura* query to *D. melanogaster* proteins) to generate common ids from our study and (Chen et al., 2015). To test for functional similarity of gene lists we estimated an odds ratio and p-value using the *fisher.test* function in R (see Zenodo archive for details). Note that our study was not intentionally designed to be compared with transcriptional plasticity following developmental acclimation (Chen et al., 2015) and that there are multiple factors differing between the two studies. Specifically, (Chen et al., 2015) 1) studied female *D. melanogaster* 2) applied egg-to-adult developmental acclimation, and 3) used a roughly similar, but not identical range of acclimation temperatures (13, 18, 23, 29°C). Thus, there are many biological reasons why the results of the studies might differ. However, finding statistical similarities between the data sets despite these confounding factors would strongly suggest functional similarities between short-term adult and long-term developmental acclimation.

### SUPPLEMENTAL FIGURES

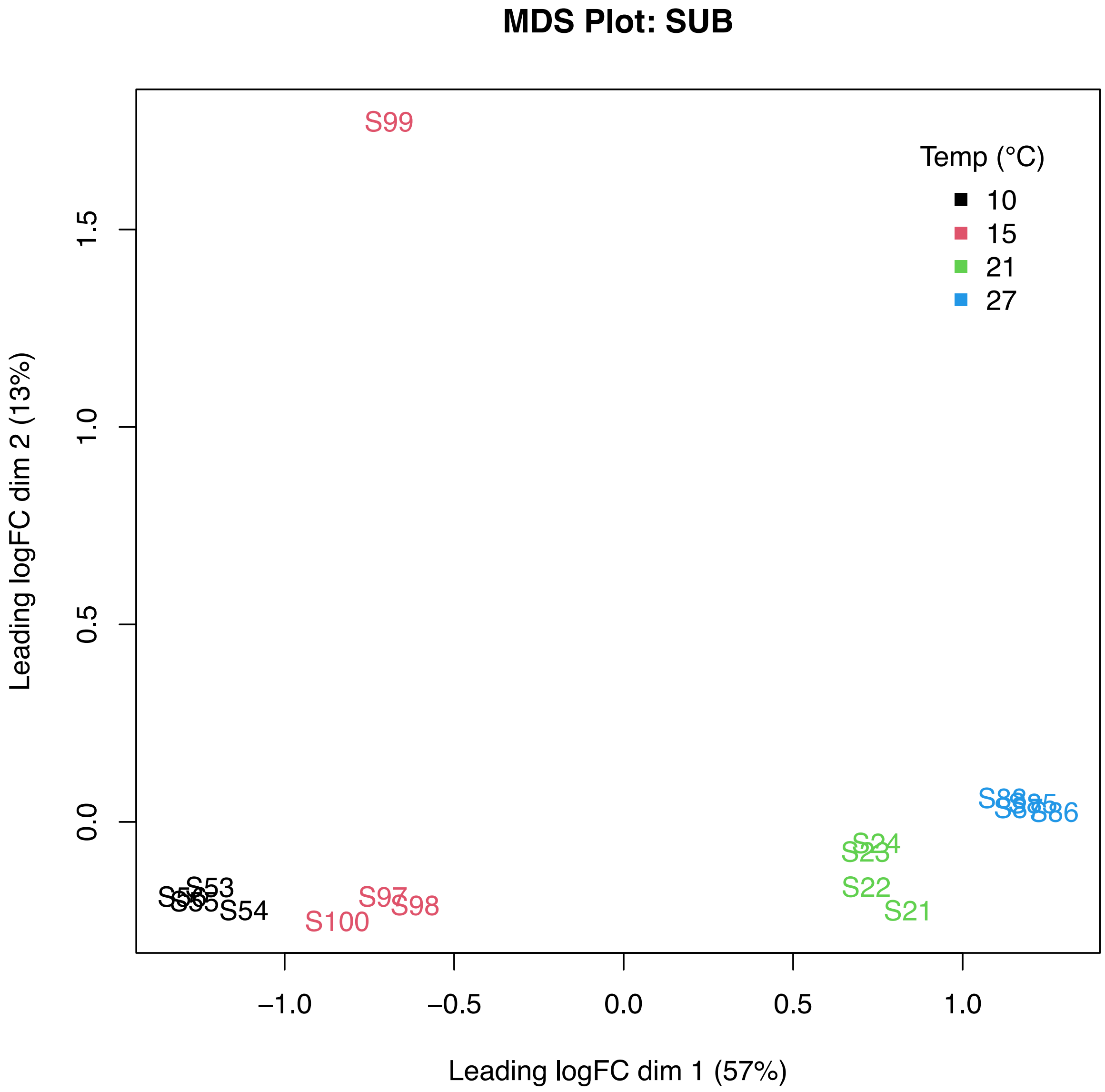


**Figure S1.** Multidimensional Scaling (MDS) analysis of the 500 most differentially abundant transcripts in adult males following acclimation at 10, 15, 21, and 27°C. Each RNA library (sample) is plotted with color indicating sampling temperature.

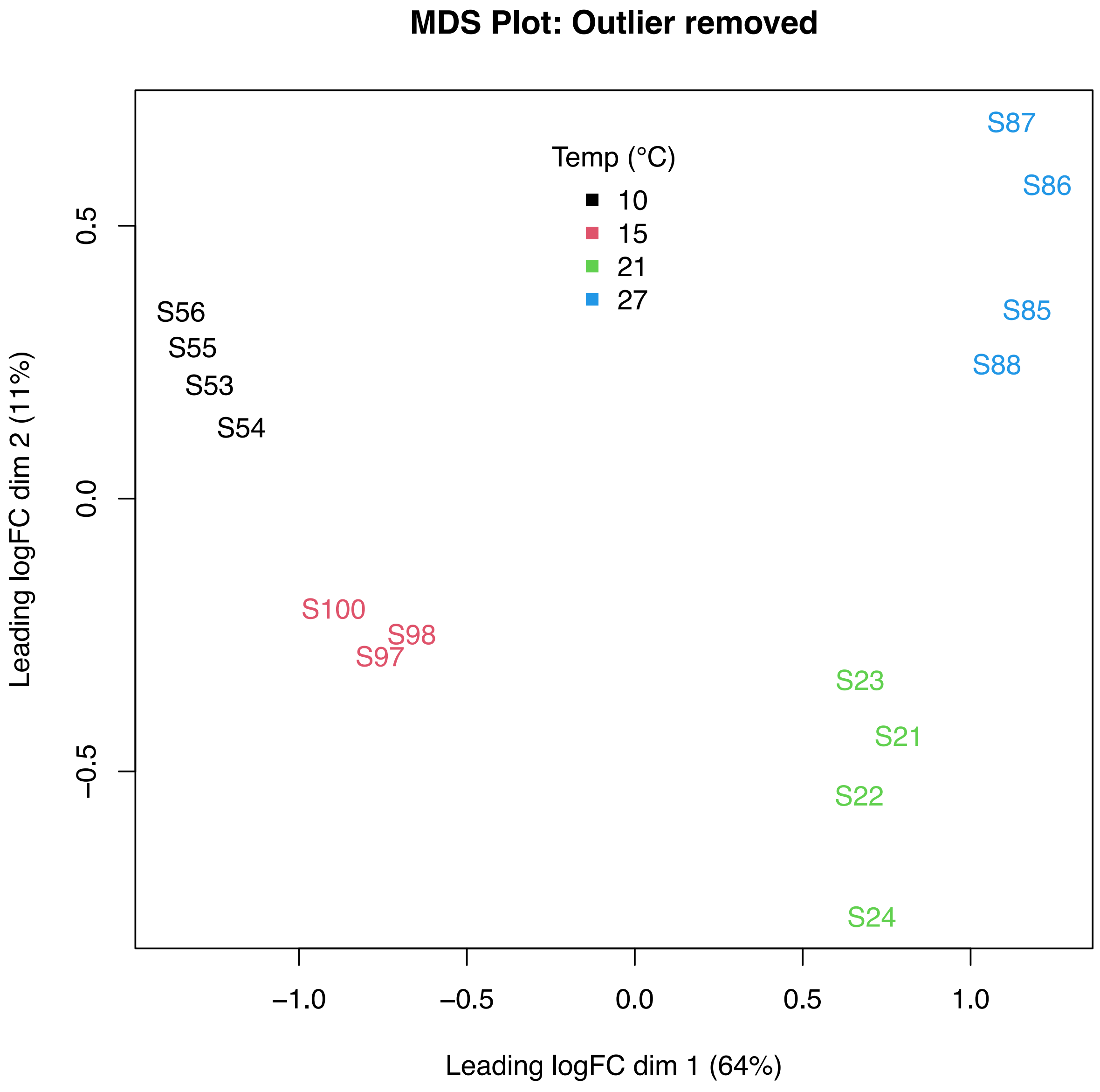


**Figure S2.** MDS analysis of the 500 most differentially abundant transcripts in adult males following acclimation at 10, 15, 21, and 27°C following removal of a female contaminated outlier sample (S99) as seen in Fig. S2.
